# Evaluate and simulate the reproduction and survival of a reintroduced population in an endangered bird species Crested ibis (*Nipponia nippon*)

**DOI:** 10.1101/2025.11.25.690618

**Authors:** Xuebo Xi, Yuansi He, Siyi Zeng, Shuai Yang, Zhuo Pan, Jia Zheng, Daiping Wang

## Abstract

1. Reintroduction programs are critical for restoring endangered species. However, scientific assessments and management recommendations based on comprehensive vital rates across the entire life cycle remain limited for such programs. The Crested ibis (*Nipponia nippon*), once nearly extinct, has been the focus of intensive conservation efforts, including reintroduction into its historical range. Here, we systematically evaluate the success of a Crested ibis reintroduction population by integrating long-term field monitoring with population dynamic modeling.
2. We conducted a systematic, multi-year (2023-2025) demographic study of a reintroduced Crested ibis population in Dongzhai National Nature Reserve, China. From 2013 to 2023, a total of 133 captive Crested ibises were released in seven batches within the nature reserve. By 2025, the wild population had exceeded 500. Based on the combination of field monitoring (n = 176 pairs) and GPS telemetry (n = 74 GPS tags, the longest monitoring time lasting 4 years), we quantified reproductive output and mortality rates across six life stages: nest building, incubation, offspring provisioning, fledgling (< 1 year), sub-adult (1-2 years), and adult (> 2 years).
3. The results show that the average clutch size of this population is 3.24 ± 0.08, and each pair of parent birds can rear 1.31 ± 0.12 nestlings to fledging each year. We found that high mortality during the incubation (32.5%, n = 425 eggs), offspring provisioning (34.1%, n = 287 chicks), and fledgling stages (38.6%, n = 63 birds). Key causes included unfertilized eggs, predation, starvation, and human disturbance. On the other hand, the mortalities in the nest building (9.1%, n = 176 pairs), sub-adult (9.8%, n = 28 birds) and adult stage (21.3%, n = 15 birds) were relatively low. Stage-structured matrix model shows a positive population growth of this reintroduced population (growth rate λ = 1.053), with the population expected to increase 13-fold over 50 years (6,858 individuals). Sensitivity analysis indicates that adult survival had the strongest influence on population growth. Correspondingly, we predict that by 2075, its distribution range will expand 13-fold in tandem with population growth, exceeding 51,000 km² and covering several surrounding cities.
4. Our comprehensive monitoring, evaluation, and simulation across the full life-cycle confirm that the reintroduced population is successfully established and self-sustaining. This finding addresses a critical research gap: the lack of a scientific and systematic framework for assessing population reintroductions in endangered species. Furthermore, we identified key conservation practices to enhance population growth and stability, including predator control, supplemental feeding, the rescue of weak chicks, and reducing human disturbance through public engagement. To ensure long-term viability, we also recommend implementing genetic management to avoid inbreeding. The resulting framework provides a practical model for improving reintroduction success and promoting species recovery in other endangered species.

## 1. Introduction

Global biodiversity is facing unprecedented threats from climate change, habitat loss, and human disturbance (Krauss et al., 2010; Urban, 2015; Pyšek et al., 2020). According to the International Union for Conservation of Nature (IUCN), over 48,600 assessed species are threatened with extinction, representing approximately 28% of all evaluated species. Effective conservation actions are urgently needed to reverse this trend. In response, numerous conservation programs have been initiated at various levels. Among these initiatives, species-oriented conservation—particularly for rare and endangered species—is highly significant (IUCN, 2025). This significance stems not only from the critically endangered status of these species, which demands immediate action, but also from their ecological and symbolic value. Specifically, such species often act as flagship or umbrella species, thereby promoting the protection of broader biodiversity and entire ecosystems (Lambeck et al., 1997; Zacharias & Roff, 2001; Qian et al., 2020). Consequently, analyzing successful case studies is essential for improving conservation efficiency and providing valuable guidance for protecting other endangered species.

Reintroduction is defined as the intentional movement and release of a species into its historical range, from which it has been extirpated, to establish a self-sustaining population (IUCN/SSC, 2013). It is a critical tool for restoring species’ historical distributions and is an essential component of modern conservation, complementing in-situ efforts by recovering species that have been lost from the wild (Armstrong & Seddon, 2008). Successful reintroductions, as documented in numerous cases, depend on integrating habitat management, predator and poaching control, and long-term monitoring. The remarkable recovery of the Arabian oryx (*Oryx leucoryx*) exemplifies this coordinated approach. After its declaration as extinct in the wild in 1972, a captive-breeding and reintroduction program launched in the 1980s—supported by habitat restoration and anti-poaching measures—enabled the wild population to surpass 1,000 individuals. This recovery prompted its downlisting on the IUCN Red List and, as a flagship species, stimulated the recovery of associated desert ecosystems (Spalton et al., 1999; Mallon et al., 2023). Similarly, the Black-footed ferret (*Mustela nigripes*) recovery underscores the necessity of holistic habitat management. Brought to near-extinction by sylvatic plague and the decline of its prairie dog (*Cynomys spp.*) prey, the ferret’s resurgence has relied on plague vaccination, prairie dog colony restoration, and pre-release conditioning. The subsequent growth of the wild population to over 300 individuals demonstrates the efficacy of securing the prey base to restore a predator population (Biggins et al., 2011; Salkeld, 2017). In contrast, the reintroduction of the California condor (*Gymnogyps californianus*) reveals a critical challenge. Although a captive-breeding program increased the population from 22 individuals in 1985 to nearly 400 by 2010, with approximately half free-flying (Walters et al., 2010), this numerical success belies a persistent vulnerability. The species’ survival remains entirely dependent on intensive management because chronic lead poisoning continues to pose a severe threat. Without a decisive reduction in lead exposure, the condor remains at high risk of re-extinction in the wild (Finkelstein et al., 2012). Collectively, these cases affirm that while reintroduction is a vital step in restoring species, identifying and mitigating core ecological and anthropogenic threats is fundamental to achieving sustainable, long-term viability.

The Crested ibis is a medium-sized wading bird distinguished by red facial skin and legs, and predominantly white plumage with cinnamon tones on the tail, abdomen, and flight feathers. Historically widespread across Northeast Asia, including China, the Korean Peninsula, Japan, and the Russian Far East (BirdLife International, 2001), its population underwent a severe decline in the mid-20th century due to habitat destruction and anthropogenic pressures such as illegal hunting and the overuse of agricultural chemicals (Ding, 2004). By the 1960s, the species was believed extinct and was consequently listed as Critically Endangered (Anderson, 1984; IUCN, 2025). However, a remnant population of just seven individuals—comprising two breeding pairs and their three offspring—was rediscovered in Yangxian County, Shaanxi Province, China, in 1981 (Liu, 1981). Following this discovery, extensive conservation efforts were implemented. These efforts have achieved remarkable success. Currently, the global population is estimated to have exceeded 10,000, and the IUCN Red List category has been downgraded to Endangered (Zheng et al., 2025). In China, reintroduction programs have expanded the species’ range beyond Shaanxi to other parts of its historical distribution, such as Dongzhai in Henan Province (Cai et al., 2024), Deqing in Zhejiang Province (Qiu et al., 2023), and Beidaihe in Hebei Province (Xie et al., 2020). Furthermore, through the provision of breeding stock from China, populations have also been established in South Korea (Lee et al., 2024) and Japan (Mochizuki et al., 2015). The recovery of the Crested ibis is a testament to decades of persistent in situ conservation and reintroduction efforts. Analyzing this successful case provides valuable insights and practical frameworks for conserving other endangered species. While existing research on the Crested ibis spans morphology, physiology (Sun et al., 2019), reproductive ecology (Ye et al., 2017), captive breeding (Okahisa et al., 2022), habitat evaluation (Li et al., 2002), and genomics (Feng et al., 2019), research on reintroduced populations remains limited. Specifically, there is a scarcity of systematic data on stage-specific survival rates and key threats across the entire life history, from the egg stage to reproductive maturity. This knowledge gap currently constrains the development of optimized conservation strategies and impedes a rigorous assessment of the population’s long-term viability.

In this study, we comprehensively assessed the survival and population viability of a reintroduced population of Crested ibis. We integrated three consecutive years of field monitoring data with five years of GPS tracking data. First, we quantified reproductive failure and stage-dependent mortality rates—specifically during the nest building, egg incubation, offspring provisioning, fledgling (<1 year), sub-adult (1–2 years), and adult (>2 years) stages—and identified their primary causes. Based on these findings, we propose targeted conservation measures to enhance breeding and survival success at each life stage. Second, using these vital rates, we simulated population dynamics over a 50-year period to evaluate whether a self-sustaining population has been established. Finally, we forecast the future distribution range of this reintroduced population. The results of this study provide a critical evaluation of the reintroduction’s success and offer a valuable framework for informing the conservation of other reintroduced populations of the Crested ibis and other endangered species.

## 2. Method

### 2.1 Study species and the study site

The Crested ibis is a long-lived, monogamous bird species that typically reaches sexual maturity between two and four years of age, with a maximum recorded reproductive lifespan of 15 years (Yu et al., 2007). Its breeding season extends from late February to late June. Both parents participate in nest building, incubation, and chick-rearing (Zhai et al., 2001). They usually build nests in tall arbors such as Masson pine (*Pinus massoniana*) and Chinese wingnuts (*Pterocarya stenoptera*). The average height of the nest-building trees is 16.1 meters with the height of the nest is 12.6 meters (Zhu et al., 2023). The species generally produces one clutch per year, consisting of 2-4 eggs (Shi et al., 1999). The incubation period is approximately 28 days, followed by a chick-rearing period of 40-45 days. After fledging, juveniles continue to depend on their parents for a period to learn essential survival skills like foraging (Ding, 2004).

The Dongzhai National Nature Reserve (31°58’ N, 114°17’ E) is a 46,800-hectare protected area situated in Luoshan County, Henan Province, on the northern foothills of the Dabie Mountains. Established to protect rare forest-dwelling birds and their habitats (Zhu et al., 2023), the reserve lies in a transitional zone between northern subtropical and warm-temperate climates. This region features a warm, humid climate and comprises well-preserved forest ecosystems interspersed with farmlands, including essential rice paddy foraging habitats for the Crested ibis. As a former part of the species’ historical range (Liu et al., 2008), Dongzhai was selected as a pioneering reintroduction site. A captive breeding program began in 2007 with the introduction of 17 individuals. Wild releases commenced in 2013, and by the final release event in 2023, a total of 133 individuals had been released into the wild across seven separate cohorts (Huang et al., 2024). The extensive forest cover, combined with paddy wetlands within the reserve, provides ideal habitat conditions for wild Crested ibis populations. This population has been continuously increasing. It is estimated that the wild population has exceeded 500. From 2023 to 2025, we continuously monitored the Crested ibis in the Dongzhai Nature Reserve and surrounding distribution areas, covering approximately 73,000 hectares.

### 2.2 Reproductive monitoring

Regular monitoring of breeding began since the first cohort of birds was released, including some newborn individuals equipped with satellite transmitters started in 2021 (n = 12). However, systematic and intensive monitoring has been carried out since 2023. During each breeding season (February to June) from 2023 to 2025, we conducted continuous observations and recorded all breeding attempts by paired Crested ibises within the study area. Data collection included nest site location, egg-laying date, clutch size, and the number of eggs hatched and chicks fledged. Routine nest checks were performed at intervals of 1-3 days to confirm the survival status of eggs or nestlings and to identify any causes of mortality. When nestlings were approximately 25-30 days old, all individuals were fitted with leg bands for identification, and a subset was fitted with satellite tracking devices (HQBG3621L model GPS tags, Hunan Global Messenger Technology Co., Ltd., Changsha, China) for subsequent monitoring. The GPS tags can record and transmit data at least once per hour, including position, temperature, acceleration, and other relevant metrics.

### 2.3 Failure/mortality and its main causing factors survey

Specifically, we conducted a comprehensive investigation of stage-specific failure and mortality (rates) in this reintroduced population, identified the primary causes of these failures, and proposed corresponding management measures to enhance population viability as follows.

#### a) attempt failures at the nest-building stage

In the early breeding season, we search for and confirm potential nesting sites or nests under construction using information such as Crested ibis calls, signs of their activity, and environmental suitability. Next, we conduct continuous monitoring of these sites or nests. Because adults are particularly sensitive to disturbance during nest building, observations were made every 3-5 days from a distance using binoculars. A nesting attempt was classified as failed when no nesting activity was detected on three consecutive visits (e.g., no additional nest material, absence of adults in the nesting area, or no new droppings beneath the nest). We have monitored 176 nesting attempts over three years (45 in 2023, 64 in 2024, and 67 in 2025).

#### b) hatching failures at the incubation stage

The incubation period was defined as beginning on the day when an adult was first observed sitting on the nest for an extended duration. Monitoring was conducted every 2-3 days, then increased to daily or every other day near the expected hatching date (the incubation period lasts approximately 28-30 days) to accurately determine hatching time. To avoid flushing incubating birds, we recorded the behavior of adults and any evidence of damaged eggs below the nest from a distance using binoculars. Hatching failures were classified into five categories (causing factors) based on the condition of the adults, incubation duration, and egg characteristics: 1) early egg loss, 2) unfertilized or rotten eggs, 3) egg predation, 4) nest desertion, and 5) extreme weather events (e.g., strong winds). The classification details and criteria are summarized in Table S1. At this stage, we monitored a total of 160 nests, including 41 in 2023, 58 in 2024, and 61 in 2025.

#### c) mortality at the offspring provision stage

During the offspring-provision period (including the offspring brood), nests were inspected every 1 to 2 days to obtain timely information on nestling status. For any dead chicks, the cause of death was inferred from field observations and necropsy examinations. Mortality causes were categorized into six groups: 1) nest predation; 2) starvation; 3) poisoning associated with the use of fertilizers or pesticides in foraging areas; 4) extreme weather events (e.g., sudden temperature drops or heavy rainfall); 5) parasitic infection, and 6) other unknown causes. The classification details and criteria are summarized in Table S2. At this stage, we monitored a total of 123 nests, including 32 in 2023, 47 in 2024, and 44 in 2025.

#### d) mortality at the fledgling stage (< 1 year)

Crested ibises typically reach sexual maturity at two years of age (Ding, 2004). We classified individuals (after fledging) of this population into three life stages: fledglings (<1 year), sub-adults (1-2 years), and adults (≥2 years). From 2021 to 2025, 74 nestlings were fitted with GPS tags. After excluding those that died prior to fledging or experienced transmitter failure, a total of 63 nestlings successfully fledged (11 in 2021, 12 in 2023, 17 in 2024, and 23 in 2025). By integrating telemetry data with field observations, we accurately determined the timing of mortality events, enabling us to estimate mortality rates for each life stage. These individuals were monitored through the fledglings stage, yielding 457 bird-months of data by September 2025. During this period, 22 fledglings died, primarily resulting from two causes: predation (by snakes and mammals) and starvation.

#### e) mortality at the sub-adult stage (1-2 years)

Excluding individuals with equipment failures and those younger than 1 year old, a total of 28 transmitter-fitted individuals were included in the monitoring of the sub-adult stage (7 in 2021, 10 in 2023, and 11 in 2024). The cumulative monitoring efforts amounted to 234 bird-months.Among them, two sub-adults died, and the cause of death was predation.

#### f) mortality at the adult stage (> 2 years)

A total of 15 individuals entered the adult monitoring stage, including 7 in 2021 and 8 in 2023. The cumulative monitoring effort reached 152 bird-months, during which three individuals died. To maximize the use of monitoring data at each stage (e.g., individuals with monitoring periods less than a full year due to equipment failures or transmitter attachment batches), we set the survival interval to one month. We used Known-fate models in Program MARK (White & Burnham, 1999) to estimate the monthly survival rate, and then calculated the corresponding annual survival and mortality rates. A detailed description of the methods and results is provided in the supplementary materials.

### 2.4 Population growth simulation

We estimated the stochastic population growth rate by simulation using a stage-based matrix (Caswell, 2006). The population was modeled with an annual post-breeding census and a 1:1 sex ratio. All adults were assumed to first reproduce at age year three and thereafter breed annually. We defined the demographic matrix as:

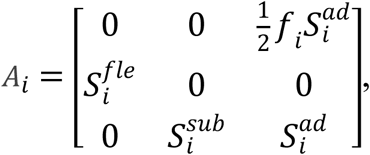

and the population size at time i+1 equals: 𝑁_𝑖+1_ = 𝐴_𝑖_ · 𝑁_𝑖_.

Each column of the demographic matrix manifests one age class. The elements of the first row represent the reproductive output produced by each age class for the next year’s population. The number of female offspring produced by each female adult that survives to breed in the next year equals to 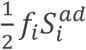. Parameter 𝑓*_i_* s the average number of fledglings produced by each female adult in year i; 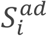 is the adult survival rate of year i. Survival coefficients over age classes are presented along the sub-diagonal of the matrix. 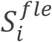 is the first year (fledglings) survival rate during year i; 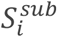 is the second year survival rate during year i.

According to the results of the autumn synchronized population survey at nocturnal roost-sites of Crested ibis in 2025 by the Nature Reserve Administration, the number of wild Crested ibis in Dongzhai is conservatively estimated to have exceeded 500. Therefore, the matrix model was initialized with a population of 500 individuals and a stable stage structure. We simulated a 50-year population dynamic using empirical parameters of reproduction and mortality across life stages. To incorporate temporal variability, we introduced stochasticity into the reproductive output at each discrete time step. Specifically, clutch size and the failure rates for the nest-building, incubation, and offspring-provisioning stages were randomly drawn from their respective 95% confidence intervals(CIs). The average number of fledglings produced per female (fᵢ) was then calculated as the product of the stochastic clutch size and the cumulative success rates of these three sequential stages (where success rate = 1 − failure rate). The same stochastic treatment was applied to the post-fledging mortality rates for the fledgling, sub-adult, and adult stages. We performed 1,000 stochastic simulations to generate the mean population trajectory and its 95% confidence interval. As the simulation tracked only the female population, the results were multiplied by two to estimate the total population size. The sensitivity and elasticity of the asymptotic population growth rate (λ) were analyzed using a population matrix projection model and its eigenvectors (Caswell, 2006). Additionally, a classic Leslie matrix was constructed based on the following life-history assumptions: a minimum reproductive age of 3 years, a maximum reproductive age of 15 years, and a maximum lifespan of 17 years. From this matrix, we derived the proportion of each age class within the stable age distribution and visualized it using an age pyramid.

### 2.5 Future distribution simulation

Based on projected population sizes, we simulated changes in the population’s distribution range over the next 50 years. The methodology consisted of three steps. First, we delineated the current distribution by constructing a minimum convex polygon around all breeding nest sites recorded in 2025, which yielded a total area of 3,700 km². Second, we calculated the number of breeding pairs in the initial 2025 population of 500 individuals by applying the stable age structure proportions derived from the demographic matrix (Section 2.4), resulting in an estimated 139 pairs. From this, we derived a mean breeding territory requirement of 26.62 km² per pair. Third, we extracted the projected number of breeding pairs from the population model at 10-year intervals from 2025 to 2075. The future distribution range at each time point was calculated by multiplying the number of pairs by the mean territory requirement (26.62 km²/pair). For cartographic representation, each resulting range was visualized as a circle of equivalent area, centered on the geometric centroid of the current distribution.

## 3. Result

### 3.1 Failures/mortality in each stage of the life history

Over three consecutive breeding seasons from 2023 to 2025, a total of 176 breeding attempts were monitored, among which 16 attempts failed (9.1%, Table 1 & Figure 1). For 32 nests located a substantial distance from the research base, observational frequency was insufficient to reliably determine egg fate. To avoid potential biases, data from these nests were excluded from analyses of the incubation and offspring provisioning stages. Among the remaining 144 breeding attempts, 131 nest-building attempts were successful, yielding a nest-building failure rate of 9.0%. These nests produced a total of 425 eggs, from which 287 chicks hatched. Eventually, 189 of these chicks fledged (Table 1). The mean clutch size was 3.24 ± 0.08 (Mean ± SE; hereafter the same), and the mean number of fledglings produced per breeding pair was 1.31 ± 0.12 (Table 1). A detailed summary of reproductive performance and stage-specific failure rates is provided in Table 1 and Figure 1.

**Figure 1.**
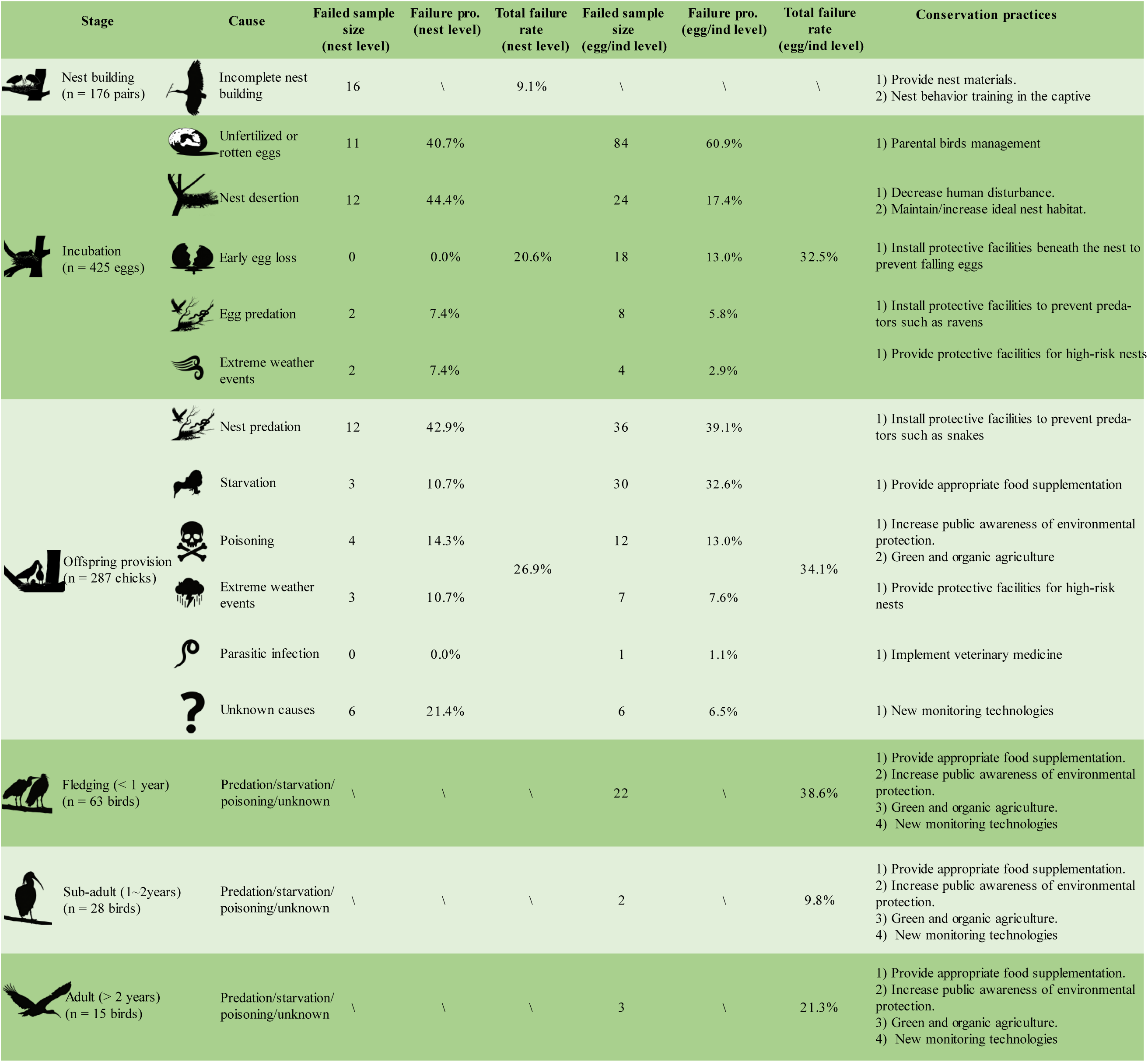
Mortality rate of the Crested ibis at each life history stage and targeted conservation strategy. Failed sample size: Number of nests or eggs that failed due to various causes. Failure pro.: Contribution ratio of each cause to the total number of failures in the current stage. Total failure rate: Overall failure rate in the current stage.

**Table 1:** Summary of breeding parameters and vital rates for the Crested ibis in Dongzhai during 2023-2025 (Mean ± SE). *Year-productivity: the average number of fledged chicks produced per breeding pair per year.

| Year | 2023 | 2024 | 2025 | Overall |
| --- | --- | --- | --- | --- |
| Number of attempts | 45 | 64 | 67 | 176 |
| Number of nests | 41 | 58 | 61 | 160 |
| Number of eggs | 115 | 163 | 147 | 425 |
| Number of chicks | 68 | 117 | 102 | 287 |
| Number of fledglings | 34 | 91 | 64 | 189 |
| Clutch size | 3.108 ± 0.121 | 3.327 ± 0.159 | 3.267 ± 0.132 | 3.244 ± 0.082 |
| Year-productivity* | 0.829 ± 0.167 | 1.655 ± 0.230 | 1.333 ± 0.198 | 1.313 ± 0.122 |
| Nest building failure rate | 0.089 ± 0.044 | 0.094 ± 0.037 | 0.090 ± 0.036 | 0.091 ± 0.022 |
| Hatching failure rate | 0.409 ± 0.045 | 0.282 ± 0.035 | 0.306 ± 0.038 | 0.325 ± 0.023 |
| Nestling failure rate | 0.500 ± 0.059 | 0.222 ± 0.038 | 0.373 ± 0.047 | 0.341 ± 0.028 |
| Fledgling mortality rate | \ | \ | \ | 0.386 ± 0.040 |
| Subadult mortality rate | \ | \ | \ | 0.098 ± 0.019 |
| Adult mortality rate | \ | \ | \ | 0.213 ± 0.031 |

We found that incubation, offspring provisioning, and fledging are the three stages during which broods experienced particularly high mortality rates (hatching failure, nestling failure and fledgling mortality, Table 1). Specifically, the mortality rate at the fledgling stage is the highest, at 38.6% (Table 1). Moreover, most individual mortalities during this stage occurred in the first half of the year (see Figure S1), likely due to the vulnerability of inexperienced fledglings to predation or starvation. Mortality at the offspring-provision stage is the second-highest, at 34.1% (Table 1). The main causes of mortality at this stage are predation and starvation. Among the nestlings that died during this stage, the proportions of deaths attributable to these two factors are 39.1% and 32.6% respectively (individual level, Figure 1). During incubation, the hatching failure rate was 32.5%. The most frequent cause was unfertilized or rotten eggs, accounting for 60.9% of failures at the egg level, followed by nest desertion (17.4%), which occurred most often during early incubation (Figure 1). Details of failure/mortality for these six stages, along with the corresponding causes and proposed conservation practices, are shown in Figure 1.

### 3.2 Population growth trend

Based on the results of the population matrix projection model, the population exhibited an asymptotic growth rate (λ) of 1.053 (95% CI: [0.970; 1.148]), an intrinsic rate of increase (r) of 0.052 (95% CI: [0.042; 0.062], Figure 2c), and a positive pyramid-shaped pattern in the age structure(Figure 2d). This indicates that, under current environmental conditions, the studied population has the potential for continuous growth, with the overall population in a healthy, positive development phase (Figure 2). The population size is expected to reach 1,400 (95%CI: [1,052; 1,926]) in 20 years, and to increase by 13-fold (compared to the population size of 500 in 2025) to reach 6,858 (95%CI: [4,122; 10,607]) (Figure 2a) in 50 years, including 3,838 adults (95%CI: [2,333; 5,889]) (Figure 2b). The sensitivity and elasticity analyses of the demographic model show that the asymptotic growth rate was most sensitive to changes in adult survival (Table S5).

**Figure 2.**
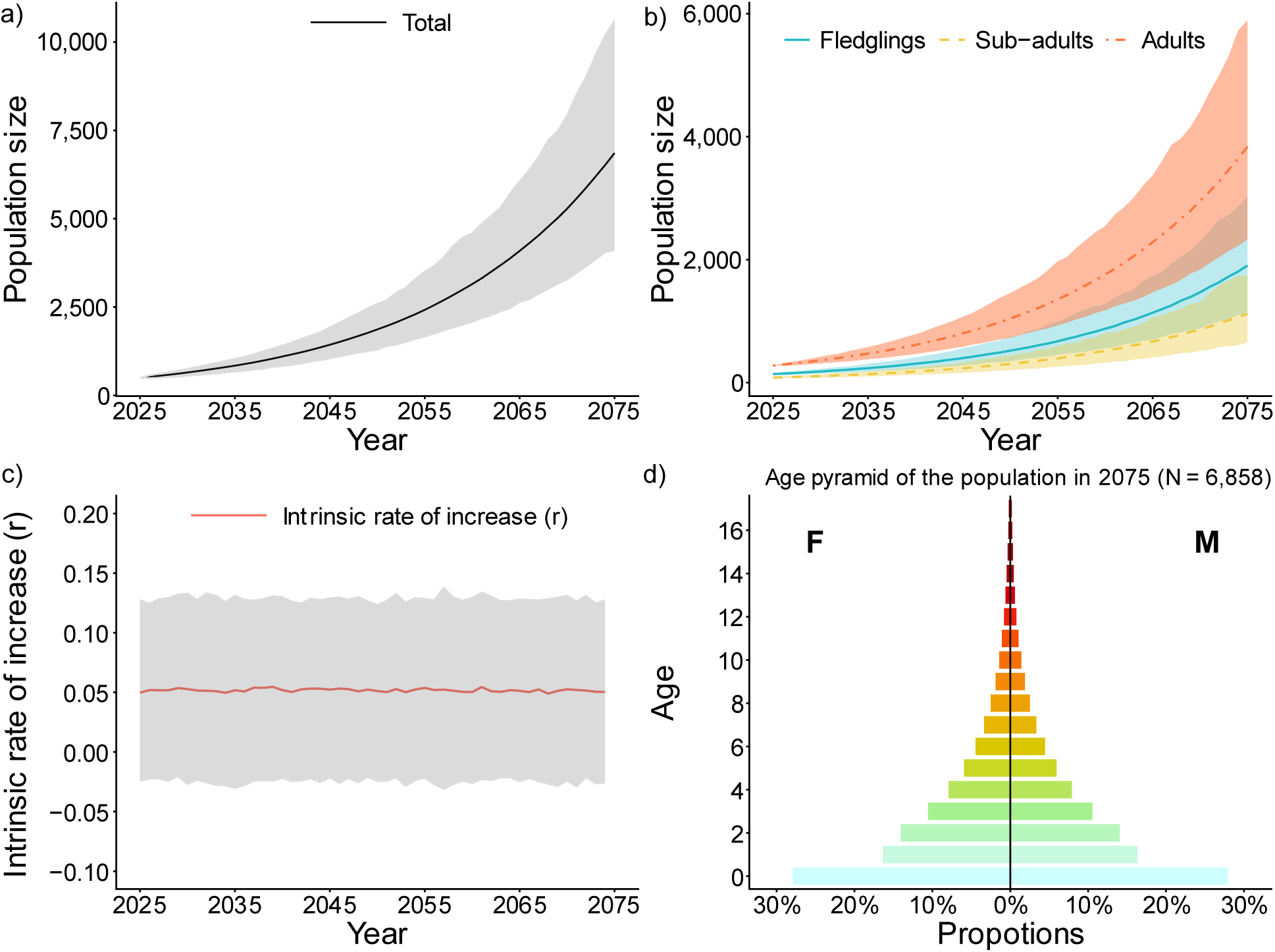
Simulation results of the population growth trend over the next 50 years. a) and b) showing the total population size and the sizes of different age categories (i.e., fledglings, sub-adults, and adults) over the next 50 years. c) the intrinsic rate of increase (r) and 95%CI of the population over the next 50 years. d) the population age-structure pyramid after 50 years in 2075.

### 3.3 Expansion of the distribution area

The area of the minimum convex polygon constructed using all the breeding nest sites in 2025 is 3,700 km², with an average area of 26.62 km² occupied per breeding pair. Based on the predictions regarding the number of breeding pairs and the expansion of the distribution range, by 2075 the radius of the distribution range of this population will increase to 3.7-fold. Correspondingly, the area will expand to 13.8-fold to reach 51,058 km², covering the surrounding Zhumadian City, Huanggang City, Wuhan City, Xiaogan City, and Suizhou City, all located around Xinyang City (Table 2 and Figure 3).

**Figure 3.**
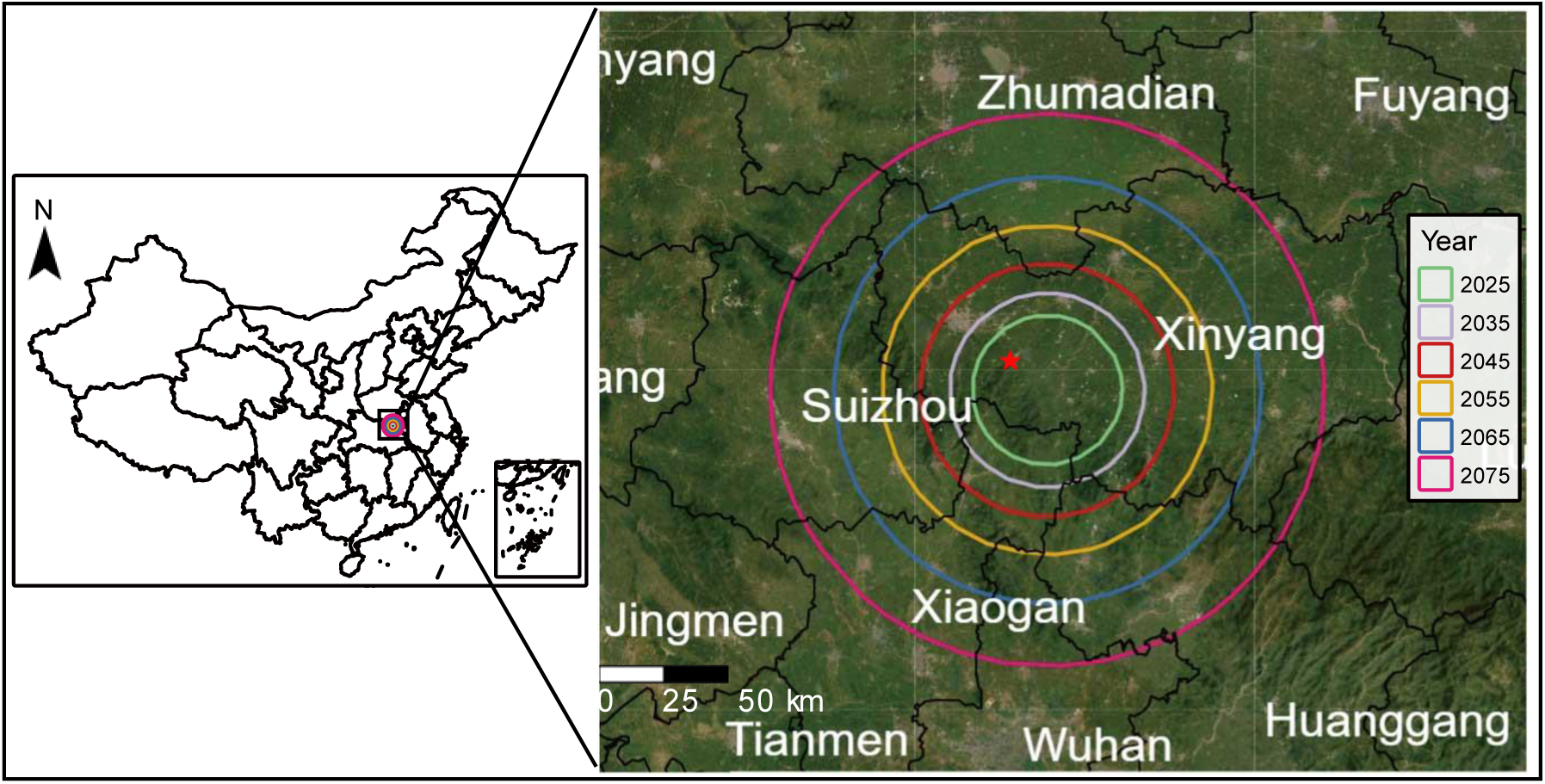
Prediction of the distribution expansion of the Crested ibis population in Dongzhai in the next 50 years. The red pentagram indicates the location of the conservation station. Six concentric circles (differentiated by color, from innermost to outermost) denote the projected distribution ranges for 2025-2075 at 10-year intervals.

**Table 2:** The number of adults, corresponding range size, and 95% confidence interval over the next 50 years.

| Year | Number of adults | Range size (km<sup>2</sup>) |
| --- | --- | --- |
| 2025 | 278 | 3700 |
| 2035 | 471 [380; 578] | 6,282 [5,058; 7,693] |
| 2045 | 802 [582; 1,070] | 10,675 [7,747; 14,242] |
| 2055 | 1,355 [930; 1,970] | 18,049 [12,378; 26,221] |
| 2065 | 2,283 [1,470; 3,364] | 30,401 [19,566; 44,776] |
| 2075 | 3,837 [2,334; 5,889] | 51,058 [31,066; 78,371] |

## 4. Discussion

In this study, we combined GPS telemetry data with intensive monitoring to systematically assess the reproduction and survival of a reintroduced Crested ibis population over three consecutive breeding seasons (2023-2025). Our study provides a detailed summary of key demographic traits across the full life cycle of this species (i.e., from nest attempting and egg laying to fledging and adulthood). Correspondingly, we identified the main factors threatening this reintroduced population and proposed pinpointed conservation practices at different stages. Furthermore, we simulated the population growth and distribution expansion in the next 50 years of this population. The results of this study provide a comprehensive picture of a successful reintroduction population. Previous studies on the Crested ibis have typically focused on one specific life stage or failed to clarify stage-specific stressors, lacking a panoramic and systematic perspective (Wang et al., 2017; Okahisa et al., 2022). Importantly, we established a research framework for reintroduction conservation initiatives in endangered species that integrates detailed monitoring, concrete conservation practices, population growth and expansion prediction, and management recommendations. This framework not only offers practical methodologies and decision support for other Crested ibis reintroduction programs but also strengthens global conservation and population recovery efforts for this flagship species. By identifying critical intervention points and refining management strategies, our work highlights the significance of linking theory and practice in endangered species reintroduction.

### Key parameters of reproduction and survival, and their main causes

The reintroduced Crested ibis population in the Dongzhai National Nature Reserve had a mean clutch size of 3.24 ± 0.08 (SE) eggs and a mean productivity of 1.31 ± 0.12 (SE) fledglings per nest per year (Table 1). Compared with the previous results from Tuohetahong et al. (2025), this productivity was lower than that of the founder population in Yangxian (1.87 ± 0.46) and another reintroduced population in Ningshan (1.76 ± 0.42). The fledgling mortality rate (0.386 ± 0.040) and the adult mortality rate (0.213 ± 0.031) were both higher than those of the Yangxian population (0.380 ± 0.095; 0.190 ± 0.040), yet lower than those of the reintroduced population in Ningshan (0.486 ± 0.038; 0.314 ± 0.034). Additionally, the sub-adults’ mortality rate (0.098 ± 0.019) was considerably lower than that of the Yangxian population (0.270 ± 0.051) and the Ningshan population (0.218 ± 0.023). To date, no other studies have shown the comprehensive, stage-specific parameters (i.e., six stages across reproduction and life-history) of reproduction and survival in this species. In this context, our investigation into the causes of reproductive failure at each stage offers valuable insights.

The nest-building stage exhibited the lowest failure rate (9.10%) among all stages. We hypothesize that failures primarily result from the parents’ inexperience with nest construction and external disturbances. As the founder populations are often captive-bred and raised in cages or with artificial nests, they lack exposure to natural nest-building processes. Therefore, we recommend that future wild-training programs incorporate training using natural nest materials and pre-constructed natural nests to improve early post-release success. Previous research for this species overlooked this stage as many begin monitoring at incubation. However, this rate is likely an underestimate due to “survivorship bias”, as early failures may have gone undetected. Meanwhile, some breeders may explore multiple potential nesting sites before selecting the ideal one. However, because some adult birds in this population have not yet been ringed, we are unable to identify breeders with multiple potential nest attempts. Nevertheless, recording breeding pairs trying to select multiple potential nest sites at the beginning and finally setting their “ideal” nest provides valuable behavioral insight. In doing so, we can evaluate the habitat quality for nest in this species (i.e., final nest site *vs*. abandoned nest sites before). In the future, as bird ringing efforts scale up, we expect to be able to more accurately estimate the failure rate at this stage and deepen our understanding of the Crested ibis habitat preference and nest-site selection.

The hatching failure rate of this reintroduced population (32.5%) is higher than that of the Ningshan (28%) and Yangxian (19.8%) populations (Yu et al., 2006; Shen, 2018). This is mainly due to the relatively high rate of infertile or rotten eggs and nest desertion rate in this population. For threatened birds, inbreeding depression is a well-documented major driver of hatching failure (Assersohn et al., 2021), and this failure rate increases with the severity of population bottlenecks (Briskie & Mackintosh, 2004). Consistent with these findings, a recent research demonstrated a negative correlation between hatching success and the inbreeding coefficient in this population (Zheng et al., 2025). Therefore, scientific pedigree management is essential for selecting specific breeding pairs for the reintroduction founder population to mitigate inbreeding. The high nest desertion rate warrants attention, accounting for 44.4% of nest failures and 17.4% of egg losses. While external disturbances are recognized as a primary cause, identifying the specific factors in the field is challenging, as nest desertion is often conflated with egg predation. Based on prior studies and our field observations, opportunistic egg predators such as the Collared crow (*Corvus torquatus*) or the Eurasian jay (*Garrulus glandarius*) may harass Crested ibises and ultimately induce nest desertion (Jiang et al., 2023). Collared crows are common and sympatric with Crested ibises in our study area. These two species were even observed nesting on the same tree. It is worth noting, however, that this issue has two sides—our field observations indicate that Crested ibises sometimes benefit from the anti-predator behavior of Collared crows, which drive raptors away from their shared nesting areas. This suggests a more complex interspecific relationship than previously assumed and warrants further research. Furthermore, human interference is another reason for nest desertion. We recorded three cases of nest desertion caused by humans—two during the Qingming Festival, when visitors entered the area for ancestral worship with firecrackers, and one during the peak of tea-picking activities. These incidents underscore the need for enhanced public conservation awareness and targeted measures to mitigate human-induced disturbances.

Predation, starvation, and pollution from excessive fertilizer and pesticide use were the primary threats across all post-hatching life stages. Snakes and raptors were the dominant predators, with the King rat-snake (*Elaphe carinata*) being the most frequently recorded nest predator. This finding is consistent with observations from the Yangxian wild population and the Ningshan reintroduced population (Yu, 2006; Jiang, 2022). Given the serious threat snakes pose to nestlings, it is necessary to enhance current protective measures by implementing additional measures. Starvation was a primary cause of mortality in Crested ibises during both the nestling and post-fledging periods. The Crested ibis is a typical asynchronously hatching species, with earlier-hatched chicks generally holding a competitive advantage. Consequently, later-hatched chicks are more likely to perish when food is limited (Ding, 2004). In reintroduction programs, providing supplemental feeding may improve breeding success (Yu et al., 2006). In combination with our results in this study, we recommend providing food supplements during years of natural scarcity. For nests with many siblings, artificially fostering, especially for later-hatched chicks, might be a good option. These interventions are crucial for enhancing reproductive output, particularly in small populations. Twelve chicks (13.04%) and two breeding adults (in 2023 and 2025) died from poisoning associated with fertilizer and pesticide use in nearby foraging areas. While modern agriculture heavily relies on agrochemicals, the promotion of organic rice cultivation in Yangxian has provided an effective example of reconciling agricultural production with habitat safety for the ibis (Wang et al., 2021).

The threat of extreme weather events to eggs and nestlings warrants serious attention. Beyond the direct impacts we observed (i.e., nests destroyed by strong winds and nestlings succumbing to severe cold and precipitation), the pathways and mechanisms through which climate affects avian reproduction are complex and often confounded with other factors. For instance, persistent heat and drought have been linked to rapid declines in nest survival for Lesser prairie-chickens (*Tympanuchus pallidicinctus*) (Grisham et al., 2016). In the context of global climate change, future research should aim to clarify the specific impact pathways of extreme weather on Crested ibis reproduction and quantify their magnitude.

Despite systematically investigating key stages that affect population viability, this study has several limitations. We could not assess the impact of inbreeding depression on mortality across these stages. Extensive research has established that inbreeding increases the homozygosity of recessive deleterious alleles, leading to abnormal embryonic development and reduced fecundity, survival, and recruitment (Brekke et al., 2010; Hoeck et al., 2015; Taylor et al., 2017; Ahlinder et al., 2021). Furthermore, individual-based models for Crested ibis population reinforcement indicate that inbreeding poses a significant extinction risk for bottlenecked populations, a critical consideration for reintroduction efforts (Zheng et al., 2025). This limitation arises from data constraints: 1) genetic samples were not collected from all individuals released initially, making pedigree reconstruction impossible; 2) the high nest locations of Crested ibises nest make capturing parents for genetic sampling prohibitively challenging; 3) a lack of systematic historical banding data of newborn individuals prevents the identification of inbreeding events. Given the essential impact of inbreeding on reproduction and survival, future research should prioritize quantifying its effects on key life-history parameters in the wild. Fortunately, remedial actions are underway to address the aforementioned constraints. Since 2023, we have systematically collected genetic samples and applied unique bands to all accessible nestlings. The first of these banded individuals began successfully reproducing in 2025, paving the way for a complete field-based pedigree database to address this knowledge gap. Additionally, the sub-adults’ mortality rate (1-2 years, 0.098 ± 0.019) was lower than that of the other two populations. Interestingly, it is also lower than the adult mortality rate in this population, which is uncommon as the adult mortality rate should be lower than that during the sub-adult stage with increasing experience. Regarding this result, we cannot rule out the possibility of sampling error. In the future, we need to further confirm these mortality estimates by increasing sample size.

### Population growth simulation

Our model projections indicate a sustained growth potential (λ = 1.053) for this reintroduced Crested ibis population, forecasting an approximately 13-fold increase over the next 50 years. Sensitivity analysis revealed that adult survival exerts the strongest influence on the population growth rate, a finding consistent with other long-lived birds, such as the White stork (*Ciconia ciconia*) and the California Condor, whose populations are also highly dependent on adult survival (Schaub et al., 2004; Walters et al., 2010). This pattern arises because adults have long reproductive lifespans, directly determining the number of breeding pairs annually. At the same time, their relatively low annual productivity (averaging 1.31 fledglings per pair) prevents rapid compensation for adult mortality. Therefore, safeguarding adult survival is the paramount strategy for sustaining population growth.

Several limitations inherent in our stage-structured matrix model should be noted. First, although the model assumed a 1:1 sex ratio, the adult sex ratio may deviate due to ecological processes like the female-biased dispersal (Hu, 2016), potentially leading to differential mortality between the sexes—a hypothesis that requires further testing. Second, the assumption of a stable age structure in 2025 may not hold, as the population has been founded through multiple releases of young birds (2-4 years old) since 2007, likely leading us to underestimate short-term reproductive output. Future models should incorporate realistic initial age structures based on banding-resighting data. Finally, the model did not account for potential future stressors, such as local habitat saturation (although individuals could freely move or dispersal in this area), disease outbreaks, or climate extremes. Thus, while current projections are optimistic, more detailed monitoring data are essential to re-evaluate population viability and enhance the robustness of our predictions.

### Distribution expansion

Our model projects that the Crested ibis population in Dongzhai will expand to a distribution range of 51,058 km² by 2075, encompassing multiple cities around Xinyang (the released city). This projection, however, relies on simplified assumptions of concentric expansion and an average home range of 26.62 km² per breeding pair. In reality, range expansion is shaped by more complex processes, primarily long-distance dispersal and metapopulation dynamics.

Sub-adults are key dispersal agents, with some undertaking long-distance movements of tens to hundreds of kilometers to colonize new habitats. Such “jump dispersal” creates isolated distribution points rather than continuous expansion. For instance, our field monitoring records document several Crested ibises dispersing nearly 70 km from the natal site, where they successfully settled and reproduced. Similarly, Tuohetahong et al. (2025) reported that a captive-bred Crested ibis released in the Yellow River Delta of Shandong completed a long-distance migration of 260 km over 11 hours. Beyond accelerating range expansion, this process alleviates density-dependent pressures in the core area, freeing up capacity for further population growth. As the range expands, the core Dongzhai population may form a metapopulation network with satellite populations established through reintroduction or dispersal. Genetic exchange and demographic rescue effects through this network would enhance overall stability—a process not captured by our current model. If suitable habitat patches exist, the actual expansion rate may thus exceed predictions.

Looking ahead, the complexity and difficulty of population management will also increase accordingly. Conservation efforts need to focus not only on the survival of individuals and the assessment of the suitability of new habitats, but also on coordinated cross-regional conservation. Establishing process-based habitat conservation goals within the species expansion range (Runge et al., 2014), identifying important ecological corridors, enhancing connectivity among populations, and promoting the formation of a metapopulation network play an important role in reducing the extinction risk of endangered species populations (Haines et al., 2006; Yang et al., 2016).

### Reintroduction evaluation

In the context of severe global biodiversity loss, the reintroduction of endangered species has emerged as a pivotal tool for species restoration. The success of these reintroductions extends beyond ensuring the persistence of the target species. As the flagship species, they can generate a significant “umbrella effect”, thereby profoundly influencing the conservation of entire ecosystems (Crespo-Gascón et al., 2019; Mekonnen et al., 2024). However, evaluating the success of reintroduction programs should not be limited to assessing the short-term survival of released individuals. Seddon (1999) posited that a successful reintroduction should sequentially achieve three progressive objectives: the survival of released individuals, the successful reproduction and rearing of offspring, and the long-term self-sustainability of the population. Robert et al. (2015) argued that reliable evaluations of ultimate success demand that populations have entered their regulatory phase. A population is regarded as self-sustaining if its long-term growth rate is positive and the probability of its persistence remains high, even in the face of environmental or demographic stochasticity (Schaub et al., 2004, 2009).

Although multiple populations have been established through reintroductions for many endangered species, the assessment of most programs remains limited to short-term monitoring, lacking systematic analysis of population dynamics and long-term projections. This gap hinders an accurate determination of their true self-sustaining status. In this context, our study focusing on a reintroduced population of an endangered species might address this gap. By integrating long-term monitoring of vital life-history parameters with population-matrix modeling and spatial-distribution simulations, we provide a comprehensive assessment of its growth trajectory and expansion potential. Our results indicate that the population has clear potential to progress towards a self-sustaining state. Therefore, our study provides a validated assessment protocol and generalizable conservation strategy for the Crested ibis reintroduction program. Furthermore, the methodology also serves as a valuable reference for reintroducing other endangered species globally, thereby optimizing resource allocation and enhancing the scientific rigor and effectiveness of conservation practices.

## 5. Conclusion

This study provides a comprehensive, full life-cycle assessment of a reintroduced Crested ibis population with intensive field monitoring and predictive modeling. Through multi-year tracking across six key life stages, we pinpointed stage-specific bottlenecks and found that there was high mortality during the stages of incubation, offspring provisioning, and fledgling. These threats are primarily driven by predation, food scarcity, and human disturbance. These findings informed targeted interventions, including predator control, supplemental feeding, and public engagement. A stage-structured matrix model projected a positive growth rate (λ = 1.053), indicating a potential 13-fold population increase over 50 years. Concurrently, the species’ distribution is forecast to expand to over 51,000 km² by 2075. Our study establishes a transferable framework for evaluating reintroduction success, linking empirical demography with spatial projections. Our approach not only supports the Crested ibis’s ongoing recovery but also provides a validated model for strengthening the scientific basis and efficacy of global conservation for endangered species.

## Notes

### Competing Interest Statement

The authors have declared no competing interest.

### Summary of Updates

Fix the display errors of individual special symbols.

### Funding

- Institute of Zoology, Chinese Academy of Sciences: 2023IOZ0104, 2024IOZ0107 to D. Wang
- National Natural Science Foundation of China: General Program, Grant No. 32570596

