## Supplemental files for "Evaluate and simulate the reproduction and survival of a reintroduced population in an endangered bird species Crested ibis (*Nipponia nippon*)"

### Supplementary material

**Calculation methods for the mortality rates of the three stages after fledging**

First, we compiled encounter history data for each individual separately to support subsequent modeling. Then, we developed a set of candidate models to simulate survival probabilities, including scenarios with constant survival and linear or quadratic changes over time. For each analysis, we compared these candidate models using Akaike's Information Criterion with a second-order correction for sample size (AICc; Burnham et al., 2011). We regarded the best-fit models as those with lower ΔAICc, higher Akaike weights ($w_i$), and fewer parameters (Burnham et al., 2011). Detailed evaluation indicators for each model can be found in Table S3. Based on the monthly survival rate estimates provided by the best-fit model, we calculated the annual survival rates for each stage and their 95% confidence intervals using the delta method (see Table S4).

**Tables and figures**

Table S1: Classification and descriptions of the causes of egg failure during the incubation stage.

| Failure cause | Incubation period | Parent bird status | Egg status |
|---|---|---|---|
| Unfertilized or rotten eggs | Middle to late stage | Normal incubation | Content deterioration or embryonic development termination |
| Nest desertion | Entire period | Incubation terminated | Eggs intact in the nest |
| Early egg loss | Early period | Normal incubation | No developmental traces |
| Egg predation | Entire period | Incubation terminated | Visible peck marks on eggshells or egg loss in the nest |
| Extreme weather events | Entire period | Normal incubation or incubation terminated | Individual or all eggs found on the ground after strong winds |

Table S2: Classification and descriptions of chick mortality causes.

| Failure cause | Offspring provision period | Number of dead chicks | Chicks' status |
|---|---|---|---|
| Nest predation | Entire period | Entire nest | Dead with wounds or entire brood missing |
| Starvation | Early to middle stage | Individual chicks | They showed severe developmental delay compared to nest-mates and had no food contents in their stomachs. |
| Poisoning | Entire period | Individual or the entire nest | Sudden death with internal signs of poisoning (hepatic/renal). |
| Extreme weather events | Early stage | Individual or the entire nest | Chicks died following adverse weather: either suddenly in the nest without trauma after cold rain, or on the ground with injuries consistent with a fall after high winds. |
| Parasitic infection | Entire period | Individual chicks | Maldevelopment was associated with a substantial parasitic load in the gastrointestinal tract identified upon dissection. |
| Unknown causes | Entire period | Individual chicks | Cause of death undetermined: advanced carcass decomposition or complete carcass loss. |

Table S3. Model selection for the survival rate (S) of fledglings/sub-adults/adults in the Crested ibis population in Dongzhai (ranked by Corrected Akaike's Information Criterion (AICc). wi: Akaike weight, K: number of parameters).

| Cohorts | Model* | K | AICc | Δ AICc | wi | Deviance |
|---|---|---|---|---|---|---|
| Fledglings | S(*tt*) | 3 | 160.803 | 0 | 0.995 | 13.633 |
| | S(*t*) | 2 | 171.27 | 10.467 | 0.005 | 26.126 |
| | S(.) | 1 | 178.413 | 17.61 | 0 | 35.287 |
| Sub-adults | S(.) | 1 | 25.049 | 0 | 0.507 | 6.922 |
| | S(*tt*) | 3 | 26.191 | 1.142 | 0.286 | 3.977 |
| | S(*t*) | 2 | 26.837 | 1.788 | 0.207 | 6.675 |
| Adults | S(.) | 1 | 31.519 | 0 | 0.637 | 9.532 |
| | S(*t*) | 2 | 33.317 | 1.798 | 0.259 | 9.277 |
| | S(*tt*) | 3 | 35.134 | 3.616 | 0.104 | 9.012 |

* Monthly survival rates were modeled as constant over time (.) or as a function of the months since entering the current stage group (either the linear effect denoted *t* or the quadratic effect denoted *tt*).

Table S4. The survival rate (S) and the 95% confidence intervals of the Crested ibis population in Dongzhai.

| Cohorts | S | SE | LCI-UCI (with 95%CI) | n |
|---|---|---|---|---|
| Fledglings | 0.614 | 0.04 | 0.541-0.697 | 63 |
| Sub-adults | 0.902 | 0.019 | 0.866-0.940 | 28 |
| Adults | 0.787 | 0.031 | 0.728-0.851 | 15 |

Table S5. Sensitivities and elasticities of lambda to underlying parameters of stage-specific fecundity and survival probability.

| Parameters | Sensitivity | Elasticity |
|---|---|---|
| Fledgling survival | 0.288 | 0.168 |
| Subadult survival | 0.196 | 0.168 |
| Adult survival | 0.664 | 0.496 |
| Year-productivity | 0.332 | 0.168 |

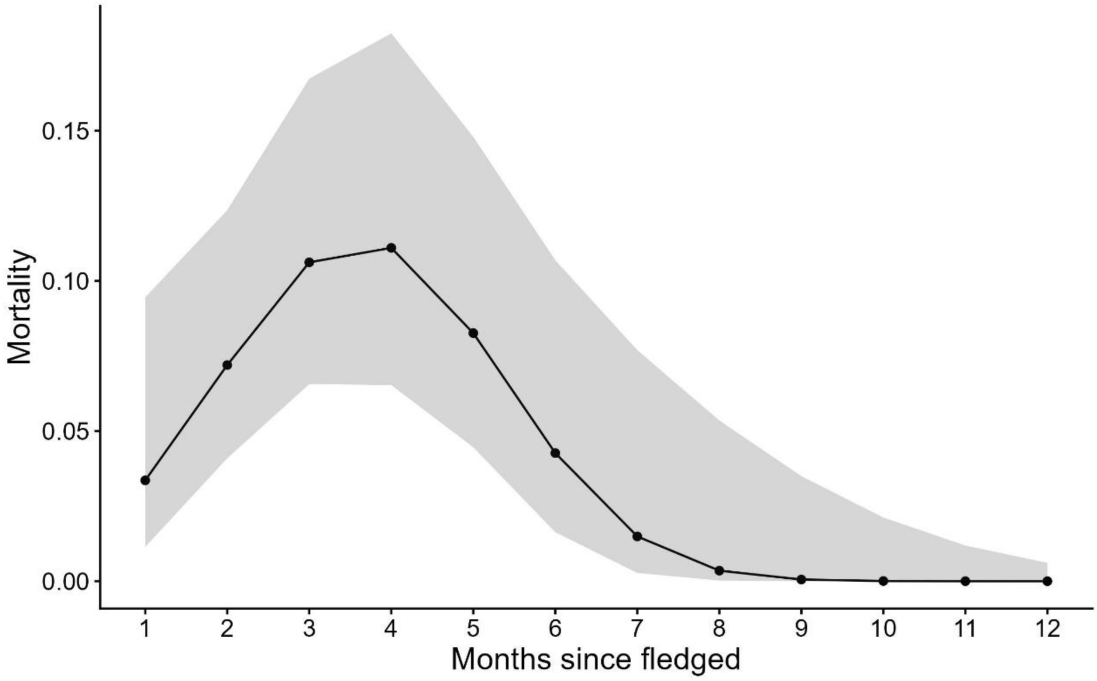


Figure S1. The monthly mortality rate of fledglings within 12 months after fledging and its 95% confidence intervals, estimated based on Known-fate model in Mark.
